# SUBTHALAMIC HIGH-FREQUENCY DEEP BRAIN STIMULATION REDUCES ADDICTION-LIKE ALCOHOL USE AND SUPPRESSES THE OVERCONSUMPTION INDUCED BY THE PEER’S PRESENCE

**DOI:** 10.1101/2023.11.21.567994

**Authors:** Lucie Vignal, Cassandre Vielle, Maya Williams, Nicolas Maurice, Mickael Degoulet, Christelle Baunez

## Abstract

**Rational:** The immediate social context significantly influences alcohol consumption in humans. Recent studies have revealed that peer presence could modulate drugs use in rats. The most efficient condition to reduce cocaine intake is the presence of a stranger peer, naive to drugs. Deep brain stimulation (DBS) of the Subthalamic Nucleus (STN), which was shown to have beneficial effects on addiction to cocaine or alcohol, also modulate the protective influence of peer’s presence on cocaine use.

**Objectives:** This study aimed to: 1) explore how the presence of an alcohol-naive stranger peer affects recreational and escalated alcohol intake, and 2) assess the involvement of STN on alcohol use and in the modulation induced by the presence of an alcohol-naïve stranger peer.

**Methods:** Rats with STN DBS and control animals self-administered 10% (v/v) ethanol in presence, or absence, of an alcohol-naive stranger peer, before and after escalation of ethanol intake (observed after intermittent alcohol (20% (v/v) ethanol) access).

**Results:** Neither STN DBS nor the presence of an alcohol-naive stranger peer modulated significantly recreational alcohol intake. After the escalation procedure, STN DBS gradually reduced ethanol consumption. The presence of an alcohol-naive stranger peer increased consumption only in low drinkers, which effect was suppressed by STN DBS.

**Conclusions:** These results highlight the influence of a peer’s presence on escalated alcohol intake, and confirm the role of STN in addiction-like alcohol intake and in the social influence on drug consumption.

## Introduction

Drug consumption is a complex behavior influenced by a multitude of factors, including environmental context and social interactions. Social context has been demonstrated to play a crucial role in modulating drug intake in both human and animal models. Indeed, social factors are known to affect multiples aspects of drug use such as initiation, maintenance, attempts to quit, and relapse, potentially boosting, mediating, or reducing addictive behaviors (Bardo et al. 2013; Pelloux et al. 2019; Venniro et al. 2018). Proximal social factors, such as the presence of peers during drug consumption, have a significant influence by decreasing or increasing drug intake (Pelloux et al. 2019; Strickland and Smith 2014; 2015). The identity of the peer is also a critical element that can modulate drug consumption. For cocaine, the presence of a non-familiar (stranger) peer leads to a stronger reduction in cocaine intake than the presence of a familiar one in both rats and humans (Giorla et al. 2022). The largest reduction was observed in presence of a stranger cocaine-naive peer. A similar beneficial effect of presence of such a peer has been also shown on addictive-like (i.e. escalated) cocaine consumption in rats (Vielle et al. preprint).

However, while the presence of peers is protective against cocaine use, its effect might be opposite for alcohol. Indeed, rats chose alcohol over social interaction in a choice procedure (Augier et al. 2022; Marchant et al. 2022), and consumed more alcohol in presence of a non-naive alcohol peer, regardless of their familiarity (Sicre et al., unpublished data). These results suggest that the influence of social presence on drug intake depends on the substance used. Nevertheless, the effect of the presence of an alcohol-naive stranger has not been assessed. Furthermore, the influence of social presence has only been examined in the context of recreational alcohol use. Consequently, the modulation of recreational and escalated ethanol intake by the presence of a stranger peer, naive to alcohol remains to be explored. Over the past few years, the Subthalamic Nucleus (STN), the only glutamatergic structure of the basal ganglia, has gained increasing significance in the reward circuit, in motivational processes and in drug addiction (Baunez and Lardeux 2011). STN inactivation induced by lesions, deep brain stimulation (DBS) or optogenetic manipulations, has shown beneficial effects across various addiction criteria: motivation (Baunez et al. 2005; Lardeux and Baunez 2008; Rouaud et al. 2010; Pelloux and Baunez 2017), escalation of drug intake (Pelloux and Baunez 2017; Pelloux et al. 2018), maintenance of escalated drug intake (Vielle et al. preprint) and compulsive drug-seeking behavior (Degoulet et al. 2021). Additionally, it has been demonstrated that the STN is involved in mediating the influence of the social context on cocaine and alcohol use. Indeed, inactivation of the STN modulates the modifications caused by the social context (presence of a peer or playback of ultrasonic vocalizations) on cocaine and alcohol intake (Giorla et al. 2022; Montanari et al. 2018; Sicre et al., unpublished data, Vielle et al. preprint; 2021).

In this study, we thus examined the influence of the presence of an alcohol-naive male stranger peer on both recreational and escalated alcohol intake in male rats. Furthermore, we investigated the contribution of the STN in these processes by using high-frequency (130 Hz) DBS.

## Materials and Methods

### Animals

32 Adult Lister-Hooded rats (∽400 g, Charles River Laboratories, Saint-Germain-sur-l’Arbresle, France) were used in these behavioral experiments. Only male rats were used to avoid inter-variability of the hormonal cycle. Animals were paired housed, in Plexiglas cages with unlimited access to food and water. Temperature- and humidity-controlled environments were maintained with an inverted 12 h light/dark cycle (light -onset at 7 pm). All experiments were conducted during the dark cycle (8am-8pm). Animals were handled daily. All animal care and use were conformed to the French regulation (Decree 2013-118) and were approved by local ethic committee and the National French Agriculture Ministry (authorization #3129.01). Experimental group sizes were determined by taking into accounts the 3Rs rules to reduce the number of animals used.

### Electrode design for STN DBS

Electrodes were made of Platinium-Iridium wires coated with Teflon (75 μm). Coating was removed over 0.5 mm at the tips and two wires were inserted into a 16 mm stainless steel tubing to form an electrode. Two electrodes, separated by 4.8 mm (twice the STN laterally), were soldered to an electric connector, allowing connection with the stimulator. Electrodes (impedance=20 ± 2.25 kΩ) and connector were subsequently deeply bound, using a custom mold and dental cement. Finally, electrodes were checked with an isolated battery to avoid electrical short circuits.

### Surgery

All the animals underwent bilateral electrode implantation into the STN. They were anesthetized with ketamine (Imalgen, Merial, 100 mg/kg, *i*.*p*.) and medetomidine (Domitor, Janssen, 0.5 mg/kg, *i*.*p*.). They also received an antibiotic treatment by injection of amoxicillin (Citramox, LA, Pfizer, 100 mg/kg, *s*.*c*.) and an injection of meloxicam (Metacam, Boehringer Ingelheim, 0.5 mg/kg, *s*.*c*.) for analgesia. Animals were then placed in the stereotaxic frame (David Kopf apparatus) for the bilateral electrodes’ implantation into the STN (coordinates in mm: -3.7 AP, ± 2.4 L from bregma, -8.35 DV from skull surface (Paxinos and Watson 2014), with the incisor bar at -3.3 mm). The electrodes were maintained with a head-cap made of dental cement and screws anchored on the skull. Rats were awakened with an injection of atipamezole (Antisedan, Janssen, 0.15 mg/kg, i.m.), an antagonist of medetomidine. Rats were allowed to recover from the surgery for at least 7 days.

### Behavioral apparatus

Behavioral experiment took place in homemade self-administration chambers (60x30x35 cm) divided into two compartments by a metallic grid. The wall of one compartment per cage was equipped with two non-retractable levers and a light above each. A recessed magazine (3.8-cm high and 3.8-cm wide located 5.5 cm above the grid floor) containing a cup liquid receptacle with an 18-gauge pipe was positioned between the two levers and connected to a pump outside the chamber. A syringe filled with ethanol was positioned on the pump and connected with the 18-gauge pipe of the receptacle via a catheter. All the chambers were controlled by a custom-built interface and associated software (built and written by Y. Pelloux).

### Behavioral procedure

An overview of the behavioral procedure is illustrated in **Fig. 1**.

**Fig. 1.**
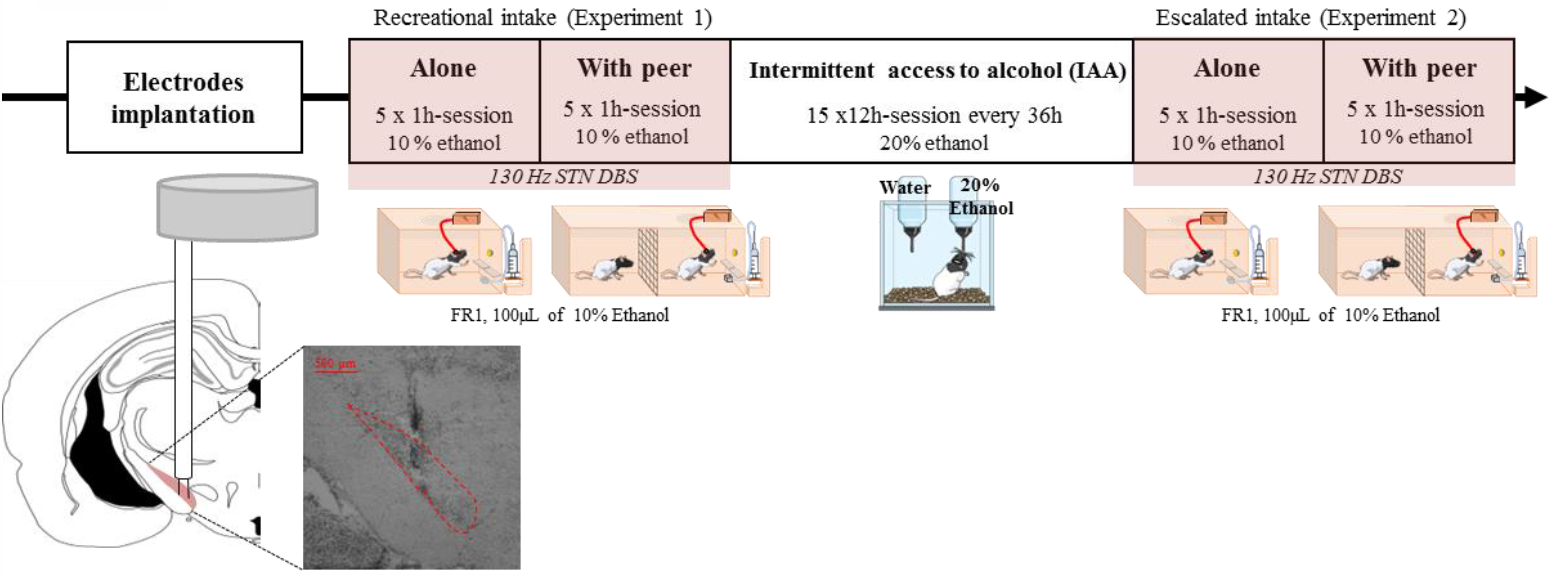
Timeline of the behavioral experiment. Top: Behavioral experiment timeline Down: (left) schematic representations of the electrode implanted in the STN, (right) Picture from a representative electrode implanted in the STN (delineated by the red dashed line)

#### 1-Experiment 1: recreational ethanol self-administration

At least one week after surgery, rats began ethanol self-administration training. The ethanol delivery was assigned to one of the two levers (‘active lever’) and counterbalanced between rats. An FR1 schedule of reinforcement was used, so each lever press delivered 100 μL of 10 % ethanol in the cup receptable and switched on the cue-light above the lever. Each delivery was followed by a 20-s time-out period, during which any press on the active lever was recorded as a perseveration but had no consequence. Presses on the other lever (‘inactive lever’) had no consequence either. Each daily session lasted 1h.

Rats were first trained to self-administer ethanol with sucrose, using a fading sucrose technique in which the sucrose concentration in the 10% ethanol solution gradually decrease over sessions until zero. They were also habituated to be connected to an electric cable (but without stimulation). Once consumption became stable (<25% of variability in the number of ethanol rewards for 5 consecutive days), animals were exposed to the testing period, during which rats were either STN-stimulated with DBS (ON group, n=16) or remained unstimulated (OFF control group, n=16). This testing period consists of 5x1h consecutive daily sessions of ethanol self-administration alone, followed by 5x1h-sessions in presence of an alcohol-naive stranger peer (‘With peer’ condition), placed on the other side of the grid. To prevent the emergence of a familiarity effect resulting from constantly using the same social stimulus, the peer was changed every day.

#### 2-Escalation of ethanol intake through intermittent access to alcohol (IAA)

After this initial testing period, animals (n=30, two animals were excluded at this point, see results section) underwent an escalation of their ethanol intake, using the intermittent access to alcohol (IAA) protocol for 15 sessions, as previously used in Pelloux and Baunez (2017). During each 12h-session (from 8:00 am to 8:00 pm), animals were isolated in a homecage with access to food and two bottles. One bottle contained water, and the other held a 20% ethanol solution. Both bottles were weighted before and after each session to calculate consumption levels. The weight of each rat was also measured prior to every IAA session. Following the 12h-period of IAA, rats were returned to their homecage with their cagemate, and observed a 36h-period without ethanol access before the next session. The placement of the bottles in the cage was alternated from session to session to avoid side preferences. There was no DBS stimulation applied during this period.

#### 3-Experiment 2: Addiction-like ethanol consumption

36 hours after the last session of IAA, rats were subjected to the same 10% ethanol self-administration procedure than for Experiment 1, for 10x1h other sessions: 5 consecutive sessions alone (‘Alone’) and 5 consecutive sessions in presence of an alcohol-naive stranger peer (‘With peer’). Operant chambers were the same as those used in the first procedure. The order of the conditions ‘Alone’ *vs*. ‘With peer’ were counterbalanced between rats. To avoid a possible long-term effect of former STN DBS (i.e., during recreational alcohol use) on addiction-like alcohol intake, we counterbalanced animals subjected to STN DBS between experiment 1 and 2. Thus, half of the animals subjected to STN DBS OFF during experiment 1 remained OFF for experiment 2, while the other half were subjected to DBS ON for experiment 2, and vice versa for the other group. The DBS was turned ON during the 10 sessions for the STN DBS group (n=15) and remained OFF for the control group (n=15).

### Deep Brain Stimulation

Electrodes were connected to homemade electric stimulation cables, themselves connected to a digital stimulator (DS8000, WPI) via a stimulus isolator (DLS100, WPI) and a rotating commutator (Plastic-One). Stimulation parameters were adapted from previous studies (Degoulet et al. 2021; Pelloux et al. 2018). Individual stimulation intensity was determined using 130 Hz frequency and 80 μs pulse width stimulation. Intensity was progressively increased until the appearance of hyperkinetic movements or abnormal behavior. Stimulation intensity (50 to 150 μA) was set just below the abnormal behavior threshold. In any case, the maximal intensity was set at 150 μA in order to prevent heating and tissue damage.

Before each behavioral session, animals were connected to the stimulation device, DBS was turned ON, and stimulation intensity was progressively increased to reach the predetermined stimulation intensity. Stimulation at 130 Hz was applied continuously during the 1-hour session.

### Histology

At the end of the experiments, rats were deeply anesthetized with an injection of pentobarbital (Euthasol Vet, Dechra, 300 mg/kg *i*.*p*.). Then, brains were extracted and rapidly frozen into liquid isopentane (-40°C) and cut in 40 μm thick frontal slices using a cryostat. A staining with cresyl violet allowed to perform histological control. Animals with incorrect electrode placement were excluded from the ON group. Representative correct electrodes implantation is illustrated in **Fig. 1**.

### Statistical Analyses

Only the number of rewards obtained (i.e., rewarded active lever presses) are shown and expressed as mean number ± SEM. Using R software, and p-value threshold set at α=0.05, data were analyzed by performing two-tailed, one, two or three-way mixed ANOVA, followed by Bonferroni post-hoc tests when applicable. Graphs were performed using GraphPad Prism 8 (version 8.0.2).

The consumption of ethanol or water in the IAA protocol was calculated as difference in bottle weight before and after each session. This difference was then divided by the weight of the rat.

In Experiment 1 and 2, the outlier’s test was conducted using the mean baseline of ethanol intake of all rats in the “alone” condition.

## Results

In total, 28 rats were included in the analyses of Experiment 1. Four animals were excluded: electrodes’ mislocation (n=2, ON group), one rat (OFF group) was identified as an outlier (estimation based on the mean baseline of ethanol intake of all rats), one rat (ON group) did not drink ethanol.

To avoid a possible long-term effect of former STN DBS during Experiment 1, we counterbalanced animals subjected to STN DBS between experiment 1 and 2. In Experiment 2 (escalated alcohol consumption), out of 30 rats, 24 were included in the analyses. 5 rats (n=2 ON and 3 OFF) did not escalate their ethanol intake in the IAA protocol, and one rat (ON group) was identified as an outlier (estimation based on the mean baseline of ethanol intake of all rats).

### Experiment 1

#### Influence of a Peer’s Presence and Contribution of STN on Recreational Alcohol Intake

To study the influence of social context and the role of STN, a three-way ANOVA was used (social context x STN DBS + session) to assess the number of rewards obtained (i.e. active lever presses outside of the timeout period). No main effects were observed for social context (F(1, 270)=0.015; p=0.901), STN DBS (F(1, 270)=0.040; p=0.842) and session (F(4, 270)=0.649; p=0.628). The interactions of STN DBS and social context were also non-significant (STN DBS x social context (F(1, 270)=0.001; p=0.972).

Indeed, in the alone condition, **control rats** (n=15) had a stable intake of ethanol *(mean 7*.*88 ± 0*.*29 rewards)*, that remained similar in presence of a peer *(mean 8*.*36 ± 0*.*48 rewards*). Thus, the presence of an alcohol-naive stranger peer, had no effect on recreational ethanol consumption.

In **STN stimulated rats** (n=13), the alcohol intake in the alone condition *(mean 7*.*60 ± 0*.*25 rewards)* was comparable to the control group. STN DBS seems not to influence recreational alcohol use in this experiment. In presence of an alcohol-naive unfamiliar peer, STN DBS did not change significantly the ethanol consumption compared to when STN-stimulated rats were in the alone condition *(mean 8*.*09 ± 0*.*15 rewards)*, and was similar to the control group’s intake (**Fig. 2a**).

**Fig. 2.**
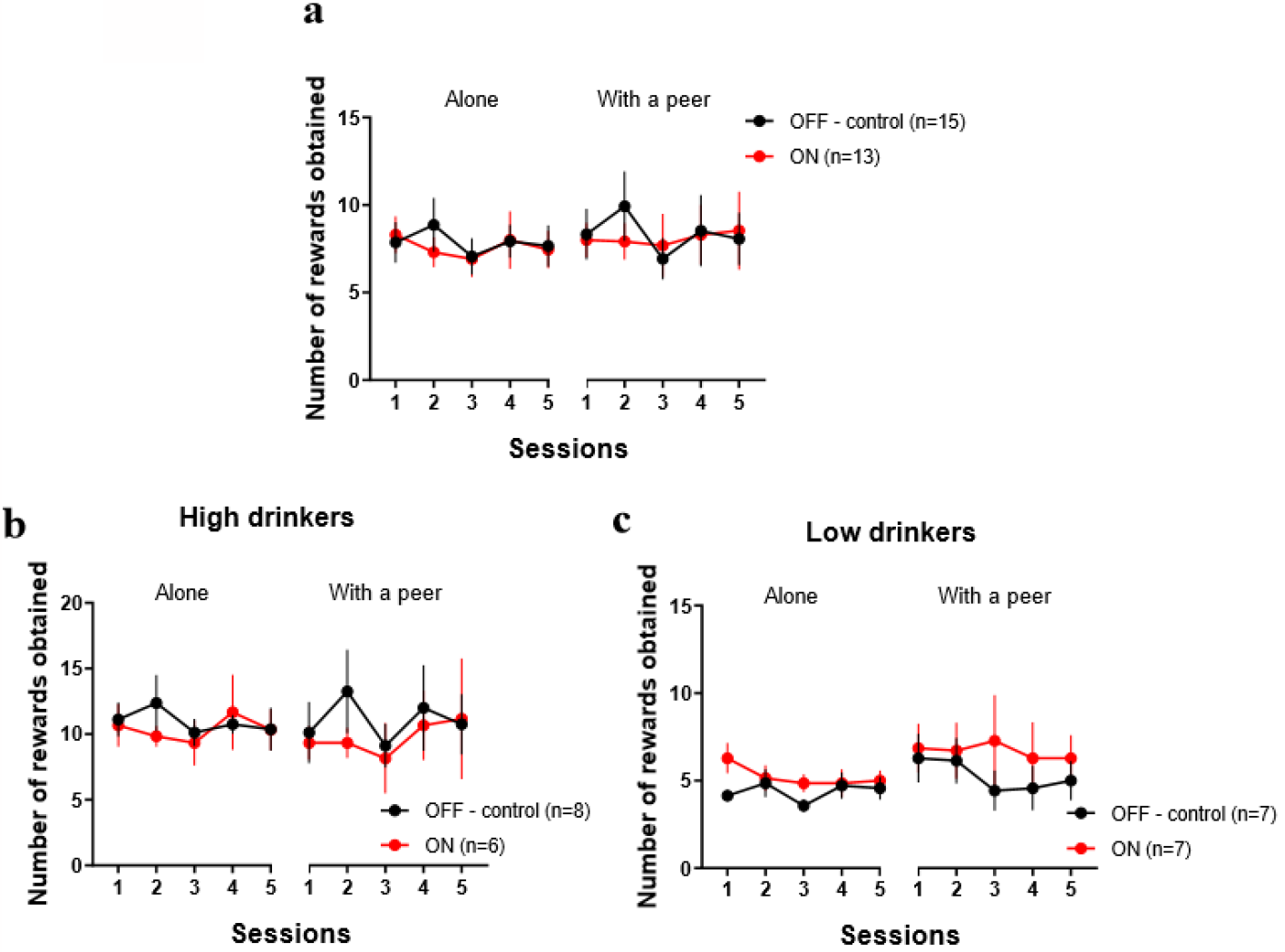
Neither peer’s presence nor STN DBS modulate recreational ethanol intake. Results are presented as mean ± SEM number of rewards obtained (i.e. number of active lever presses) per 1h-session. (a) Number of rewards obtained for non-STN-stimulated (black dots – OFF - control) and stimulated (red dots – ON) groups during five sessions in the alone condition (*7*.*88 ± 0*.*29 vs. 7*.*60 ± 0*.*25 rewards* respectively) and five sessions with an alcohol-naive stranger peer (*8*.*36 ± 0*.*48 vs. 8*.*09 ± 0*.*15 rewards* respectively) (b) Number of rewards obtained in control (black dots – OFF - control) and STN-stimulated (red dots – ON) High drinkers in the alone condition (*10*.*95 ± 0*.*40 vs. 10*.*37 ± 0*.*40 rewards* respectively) and in a presence of an alcohol-naive stranger peer (*11*.*05 ± 0*.*72 vs. 9, 73 ± 0*.*53 rewards*, respectively) (c) Number of rewards obtained in control (black dots – OFF - control) and stimulated (red dots - ON) Low drinkers in the alone condition (*4*.*37 ± 0*.*23 vs. 5*.*23 ± 0*.*27 rewards*, respectively) and in a presence of an alcohol-naive stranger peer (*5*.*29 ± 0*.*39 vs. 6*.*69 ± 0*.*19 rewards*, respectively)

### Influence of the Alcohol Preference

Because substantial variations in alcohol consumption and preference were observed between rats, they were split into two groups: high drinkers (HD) and low drinkers (LD) depending on their mean ethanol consumption in the alone condition (Spoelder et al. 2015). The effects of the social context and STN DBS were assessed with a three-way ANOVA (social context x STN DBS + sessions) independently for each group according to their preference for ethanol.

**In HD rats** (n=14, **Fig. 2b**), there were no effect of the STN DBS (F(1, 130)=0.061; p=0.805), nor of the sessions (F(4, 130)=0.759; p=0.554) nor of the interaction STN DBS x social context (F(1, 130)=0.159; p=0.691), although a trend for the social context effect (F(1, 130)=3.518; p=0.063). Indeed, HD control rats (n=8) had similar ethanol consumption in the alone condition *(mean 10*.*95 ± 0*.*36 rewards)* and in a presence of an alcohol-naive stranger peer *(mean 11*.*05 ± 0*.*72 rewards)*. STN DBS did not affect alcohol consumption in HD animals (n=6) (*alone*: *mean 10*.*37 ± 0*.*40, with a peer: mean 9*.*73 ± 0*.*53 rewards*), that was equivalent to the control group in the same circumstances.

**In LD rats** (n=14, **Fig. 2c**), the ANOVA analysis indicated a trend for an STN DBS effect (F(1, 130)=3.669; p=0.058) and for the social context (F(1, 130)=3.415; p=0.067), but no significant effect of the repeated sessions (F(4, 130)=0.459; p= 0.766), nor interaction STN DBS x social context (F(1, 130)=0.356; p=0.552).

LD control rats (n=7) consumed ethanol at the same level in the alone condition (*mean 4*.*37 ± 0*.*23 rewards)* and in presence of a peer (*mean 5*.*29 ± 0*.*39 rewards)*. STN DBS in LD rats (n=7) did not change ethanol consumption when compared to that of control animals, regardless of whether they were in the alone condition *(mean 5*.*23 ± 0*.*27 rewards)* or in a social context *(mean 6*.*69 ± 0*.*19 rewards)*.

Overall, our results showed that despite tendencies for slight modulations, neither the presence of an alcohol-naive stranger peer, nor the STN DBS alters significantly recreational alcohol consumption, regardless of alcohol preference (i.e., low *vs*. high drinkers).

### Experiment 2: Influence of peer’s presence and STN contribution following escalation of ethanol use

#### Rats subjected to IAA procedure escalated their ethanol consumption over sessions

Alcohol consumption during IAA was analyzed with a two-way ANOVA with STN DBS group as between factors (control group *vs*. “to be stimulated group” although there is no stimulation during IAA) and sessions as within factors. During the intermittent access to alcohol, rats showed a progressive increase of their alcohol consumption **(Fig. 3a**) over the sessions (F(4.34, 95.43)=18.083; p<0.0001), with no difference between the control group and the “to be stimulated” group (F(1, 22)=1.197; p=0.286) and no interaction between these two factors (F(4.34, 95.43)=1.749; p=0.140). Post-hoc analyses confirmed the progressive increase of alcohol intake over the sessions, reaching a level of consumption during the last session (*mean 3*.*01 ± 0*.*35 g/kg ethanol)* that was significantly higher than that of the first session (*mean 1*.*66 ± 0*.*13 g/kg ethanol*, Bonferroni’s test: p=0.00015).

**Fig. 3.**
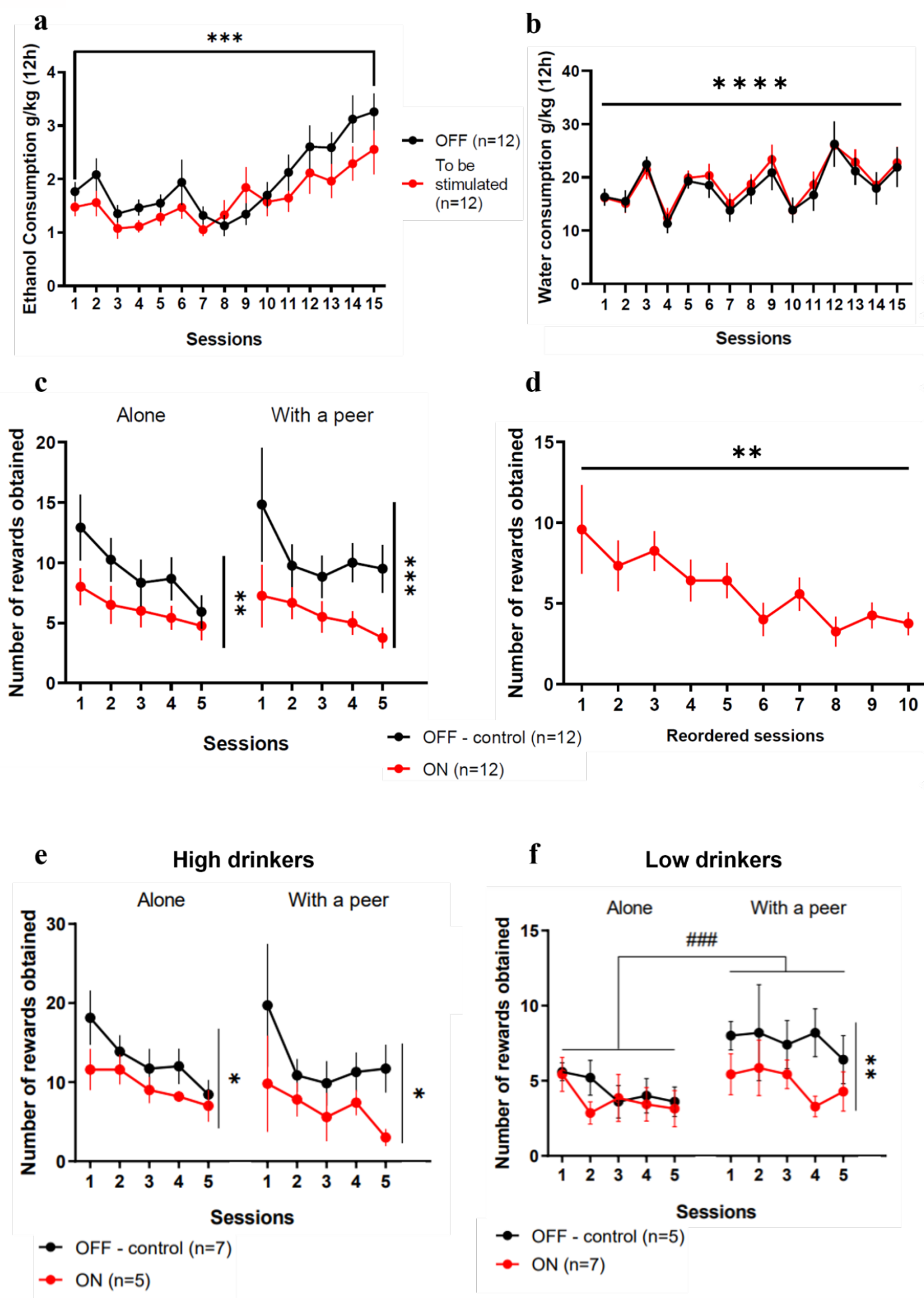
STN DBS reduces the escalated intake of alcohol and suppresses the increased consumption induced by the peer’s presence. Results are presented as mean ± SEM. (a) Ethanol intake in control (black dots - OFF) and “to be stimulated” (red dots – ON) groups across the 15 sessions of intermittent access to 20% ethanol. Both groups increased their consumption from session 1 (*1*.*66 ± 0*.*13 g/kg ethanol)* to session 15 (*3*.*01 ± 0*.*35 g/kg ethanol)*. (b) Water intake in OFF (black dots - OFF) and “to be stimulated” (red dots - ON) groups across the 15 sessions of intermittent access to 20% ethanol. Both groups consumed the same amount of water between session 1 (*16*.*22 ± 0*.*92 g/kg water)* and session 15 (*22*.*39 ± 2*.*27 g/kg water)*. (c) Number of rewards obtained after the loss of control of alcohol intake for non-stimulated (black dots – OFF - control) and STN-stimulated rats (red dots – ON) during five 1h-sessions in the alone condition (*9*.*30 ± 1*.*10 vs. 6*.*13 ± 0*.*55 rewards*, respectively) and five 1h-sessions with an alcohol-naive stranger peer (*10*.*58 ± 1*.*08 vs. 5*.*63 ± 0*.*62 rewards*, respectively) (d) Number of rewards obtained by the stimulated rats after reordering the sessions in the proper sequence irrespective of the social context. ANOVA revealed a general session effect, reflecting a gradual decrease of the ethanol intake over time. (e) Number of rewards obtained in control (black dots – OFF - control) and STN-stimulated (red dots - ON) High Drinkers after a loss of control, in the alone condition (*12*.*83 ± 1*.*59 vs. 9*.*48 ± 0*.*92 rewards* respectively) and in a presence of an alcohol-naive stranger peer (*12*.*69 ± 1*.*78 vs. 6*.*72 ± 1*.*15 rewards*, respectively) (f) Number of rewards obtained in control (black dots – OFF - control) and stimulated (red dots – ON) Low Drinkers after a loss of control, in the alone condition (*4*.*40 ± 0*.*42 vs. 3*.*74 ± 0*.*45 rewards*, respectively) and in a presence of an alcohol-naive stranger peer (*7*.*64 ± 0*.*34 vs. 4*.*86 ± 0*.*48 rewards*, respectively) *\* p<0*.*05; ** p<0*.*01; *** p<0*.*001; **** p<0*.*0001* *### p<0*.*001 between “alone” and “with peer” conditions in the non-stimulated (*OFF - control*) group*

Two-way ANOVA (STN DBS group x sessions) of water consumption during IAA (**Fig. 3b**) revealed a main effect of session (F(5.40, 118.80=7.893; p<0.0001) and no difference between the two future groups (F(1, 22)=0.170; p=0.5) nor the interaction between session x STN DBS (F(5.40, 118.80)=1.011; p=0.418). The session effect is due to inter-session variability but no difference between the first (*mean 16*.*22 ± 0*.*92 g/kg water)* and the last sessions (*mean 22*.*39 ± 2*.*27 g/kg water*, Bonferroni’s test, p=1) was found.

As a result, both control and ‘to be stimulated’ groups exhibited comparable escalation and loss of control only over alcohol intake.

#### Influence of Peer’s Presence and STN Contribution on Escalated Alcohol Intake

To assess the effect of the peer’s presence and STN DBS after a loss of control over alcohol use, a three-way ANOVA was conducted (STN DBS x social context x sessions). This investigation disclosed significant main effects of STN DBS (F(1, 215)=39.752; p < 0.0001) but no effect of the social context (F(1, 215)=0.232; p=0.631), and sessions (F(4, 215)=0.337; p=0.852). No significant interaction was found (STN DBS x social context (F(1, 215)=1.230; p=0.269; STN DBS x sessions (F(4, 215)=1.090; p=0.363; social context x session (F(4, 215)=0.073; p=0.990) and STN DBS x social context x session (F(4, 215)=0.250; p=0.909).

In the **control group** (n=12, **Fig. 3c black**), after escalation of alcohol intake, the introduction of an alcohol-naive stranger peer, did not modulate ethanol consumption; control rats displayed equivalent ethanol intake whether in the alone condition *(mean 9*.*30 ± 1*.*10 rewards)* or in presence of the peer *(mean 10*.*58 ± 1*.*08 rewards*, Bonferroni’s test, p=0.341).

When **STN DBS** (n=12, **Fig. 3c**) was applied, post-hoc analysis revealed that in the alone condition, rats consumed a significantly less ethanol (*mean 6*.*13 ± 0*.*55 rewards*, Bonferroni’s test, p=0.004) than the control group in the same alone condition. This decreased consumption persisted in the presence of a peer, where STN-stimulated rats consumed significantly less (*mean 5*.*63 ± 0*.*62 rewards*, Bonferroni’s test, p=0.0004) than the control groups in the same condition. However, within the stimulated group, no significant difference was observed between the level of consumption in the two conditions ‘alone’ and ‘with peer’ (Bonferroni’s test, p=0.567). Together, these results suggest that STN DBS reduces escalated ethanol intake, independently of the social context.

To assess a possible gradual effect of the STN DBS on ethanol intake, we reordered the sessions in the proper sequence independently of the social conditions (i.e., the order of passage in the two social conditions were counterbalanced between animals, **Fig. 3d**). We then performed a one-way ANOVA (reordered session) to assess the effects of the STN DBS over time on alcohol intake. Our results revealed a significant effect of the reordered session (F(9, 109)=2.396; p=0.008), confirming that the STN DBS progressively decreased alcohol intake.

### Impact of Alcohol preference Following a Loss of Control over Alcohol Intake

An additional possible contributor to inter-individual variation is the individual alcohol preference. This alcohol preference is calculated on the level of alcohol consumption in the alone condition after the escalation procedure. This alcohol preference allows splitting rats into two categories: High Drinkers (HD) and Low Drinkers (LD).

**For HD rats** (n=12, **Fig. 3e**), a three-way ANOVA was run to explore the influence of social context, STN DBS, and repeated sessions. While no impact of the social context (F(1, 95)=0.724, p=0.397) and sessions (F(4, 95)=0.698; p=0.595) emerged, effect of STN DBS (F(1, 95)=10.003; p=0.002) was significant. No interaction effects were found (STN DBS x social context: F(1, 95)=0.792; p=0.376), STN DBS x session: F(4, 95)=0.434; p=0.784, social context x session: F(4, 95); p=0.248; p=0.910 and STN DBS x social context x session: F(4, 95)=0.192; p=0.942) .

HD control animals (n=7) had a similar ethanol consumption in the alone condition and in the presence of a peer (*mean 12*.*83 ± 1*.*59 and 12*.*69 ± 1*.*78 rewards respectively*, Bonferroni’s test, p=0.948), revealing that the peer presence did not influence escalated alcohol use in HD rats. The STN DBS reduced the consumption in HD STN-stimulated animals (n=5) compared to the HD control group in both social conditions: alone (Bonferroni’s test, p=0.037, *mean 9*.*48 ± 0*.*92 rewards* for stimulated animals) and in presence of a peer (Bonferroni’s test, p=0.021, *mean 6*.*72 ± 1*.*15 rewards* for STN-stimulated rats). However, no significant difference was found in alcohol intake between the two social conditions within the HD STN-stimulated group (Bonferroni’s test, p=0.117), revealing the beneficial effect of the STN DBS on escalated alcohol use in HD rats, independently of the social context.

**For LD rats (**n=12, **Fig. 3f**), the three-way ANOVA revealed a significant effect of the social context (F(1, 95)=10.631; p=0.002) and STN DBS (F(1, 95)=10.108; p=0.002), but no influence of the session (F(4, 95)=0.443; p=0.778). No significant interaction was found (STN DBS x social context: F(1, 95)=2.919; p=0.091, STN DBS x session: F(4, 95)=0.395; p=0.812, social context x session: F(4, 95)=0.317; p=0.867 and STN DBS x social context x session: F(4, 95)=0.315; p=0.868).

LD control animals (n=5) consumed more alcohol in presence of a peer (*mean 7*.*64 ± 0*.*34 rewards*) than when they were in the alone condition (*mean 4*.*40 ± 0*.*42 rewards*, Bonferroni’s test, p=0.0006). Although STN DBS (n=7) did not affect ethanol intake when rats were in the alone condition (Bonferroni’s test compared to LD controls, p=0.366, *mean 3*.*74 ± 0*.*45 rewards*), it prevented the overconsumption induced by the presence of a peer (*mean 4*.*86 ± 0*.*47 rewards*) (lower consumption in LD STN DBS compared with LD controls, Bonferroni’s test, p=0.005). Consequently, STN DBS effectively suppressed the excessive consumption triggered by the presence of a peer in LD rats.

## Discussion

The present study showed that neither the presence of an alcohol-naive stranger peer, nor STN DBS, significantly modulate the recreational use of alcohol. In contrast, after a loss of control over ethanol intake, STN DBS induced a progressive decrease of the alcohol use over time. After escalation of ethanol intake, the presence of an alcohol-naive stranger peer modulated the consumption depending on the initial alcohol preference. HD rats were insensitive to the presence of the peer, and maintained a stable high level of ethanol intake, while LD rats exhibited an overconsumption in presence of the peer. STN DBS decreased ethanol intake in HD rats in both conditions (alone and with peer), while it only reduced the overconsumption induced by the peer’s presence in LD rats.

### Presence of a peer during alcohol consumption: Importance of the peer’s identity

Studies investigating the effect of a peer’s presence on alcohol self-administration have reported yielded mixed results. Several showed that the presence of a social partner drinking alcohol promotes alcohol drinking in humans (Dallas et al. 2014; Larsen et al. 2009). Our recent study in rats further revealed that the presence of a peer, whether familiar or stranger, but who had a regular consumption of alcohol, induced an increase of recreational alcohol consumption (Sicre et al., unpublished data). In the present study, the presence of an alcohol-naive stranger peer during recreational use of alcohol had no significant influence. These results highlight that the peer’s history with alcohol plays an important role. If peers consumed alcohol orally recently before being used as the “peer”, it is possible that olfactory cues of alcohol consumption could be transmitted to the subject rat through mouth-to-mouth contacts. These ethanol cues might induce a social transmission of ethanol preference, as observed for food (Strupp and Levitsky 1984), and this would result in an overconsumption of alcohol. Such olfactory cues could not be transmitted by a completely alcohol-naive peer, preventing alcohol preference facilitation and therefore no changes in consumption for the subject rat. On the other hand, the absence of increase of alcohol intake during the recreational experiment could be due to a ceiling effect that does not leave room for an increase. However, we have observed an increased effect by the presence of a peer in LD rats after escalation of alcohol consumption and the level reached then by these LD was equivalent to that recorded in the recreational experiment. So, although there was room for an increase in consumption in the recreational experiment, the peer presence had no significant effect. The ceiling effect is thus probably to be ruled out.

Once the control over ethanol intake was lost, the influence of an alcohol-naive peer varied depending on alcohol preference. For LD rats, the presence of a peer induced an overconsumption of ethanol while for HD rats, there was no modulation. An appealing hypothesis would be that after an escalation of alcohol intake, LD rats had an enhanced sensitivity to the presence of a peer. Indeed, social interactions are rewarding (Krach 2010), and after undergoing intermittent ethanol exposure during late adolescence, male rats having low doses of ethanol in adulthood displayed an increased social behavior (Varlinskaya et al. 2014). The potentiation between the rewarding effect of low doses of alcohol and social interaction could be magnified, resulting in the overconsumption even if the peer is an alcohol-naive stranger peer. In the case of HD rats after a loss of control, their lack of sensitivity to the peer’s presence could be attributed to a preference for alcohol consumption over social interaction. It has been shown that when rats had a high dose of ethanol, they show less social investigation, contact behavior and social avoidance (Varlinskaya and Spear 2002). Moreover, after a protocol of intermittent access to alcohol rats exhibit preference for alcohol over social rewards (Augier et al. 2022; Marchant et al. 2022). In our experiment, however, rats were not offered the choice, the social presence was imposed, and the rats could consume alcohol while interacting with a peer as much as they liked.

Nonetheless, from ours results, the presence of a stranger peer that never consumed ethanol had not much detrimental effect, which is in contrast to the consequences of presence of a peer that has previously consumed alcohol. These results confirm the significant influence of the immediate social context on drug consumption. This effect is influenced by factors such as the peer’s identity (e.g., familiarity, drug history) and the specific type of drug being used (e.g., psychostimulant *vs*. depressive drug).

### STN as a neurobiological substrate …

#### …Of escalated but not recreational alcohol use…

During recreational alcohol use (Experiment 1), the present study showed no significant effect of STN DBS on ethanol intake. These results are consistent with a previous work showing that lesion of STN did not affect rats’ alcohol intake in a forced consumption test, and in a two-bottle choice protocol (Lardeux and Baunez 2008). Moreover, similar to alcohol, STN manipulation was ineffective at decreasing recreational cocaine use (Baunez et al. 2005; Rouaud et al. 2010; Giorla et al. 2022; Vielle et al. preprint). However, while no effort was required in the present study that used continuous reinforcement, when the demand was increased, as in a progressive ratio task, STN manipulation was efficient at reducing cocaine motivation (Baunez et al. 2005; Rouaud et al. 2010). For alcohol, this effect has been shown to depend on the loss of control over ethanol intake. If rats did not undergo an IAA protocol, the effect of STN lesion depended on the rat’s alcohol preference (Lardeux and Baunez 2008). After an IAA protocol, STN lesion decreased the level of ethanol motivation (Pelloux and Baunez 2017). Thus, like for cocaine, STN manipulation could blunt alcohol addiction-like criteria, without altering recreational drug use.

Indeed, applying DBS at high frequency to the STN after escalation of ethanol intake reduced alcohol consumption. These results are in line with previous works showing that lesion of STN prevented escalation of ethanol intake and diminished the rebound intake after abstinence (Pelloux and Baunez 2017). This has been shown for cocaine, heroin and alcohol (Pelloux and Baunez 2017; Pelloux et al. 2018; Vielle et al. preprint; Wade et al. 2016). Interestingly, in contrast to our former results with cocaine that failed to show an effect of STN DBS on the maintained escalated intake (Pelloux et al. 2018), it was here reduced for alcohol. This disparity in the impact of STN DBS on escalated alcohol and cocaine consumption could potentially be attributed to the type of drug available. Cocaine is a psychostimulant, while alcohol has depressant effects. The road of administration could also explain the differences observed between alcohol and cocaine. Alcohol is consumed orally, leading to different pharmacokinetics and longer absorption time compared to intravenously administration that enters directly in the bloodstream (Robinson et al. 2002). This delay deferred the activation of the reward system and pleasurable effects (Robinson et al. 2002). Introducing a delay before obtaining a reward has been shown to decrease motivation (Jarmolowicz and Hudnall 2014). Consequently, it might thus be comparatively easier to reduce motivation for alcohol compared to cocaine with STN DBS. However, one might also consider that having a direct sensory input with alcohol in the mouth, but also its smell, could lead to anticipation of ethanol effects, while intravenous injection might be less efficient at anticipating the physiological and psychoactive effects of cocaine. The differences are therefore difficult to simply explain with the mean of administration. It is interesting to note however that, although the DBS or lesion procedure did not lead to changes in sustained escalated intake of cocaine, a precise targeting of the STN using optogenetics has been shown to be efficient (Vielle et al. preprint).

Finally, differences in the experimental protocol would account for such differences. Indeed, cocaine escalation, and maintenance in Pelloux et al. (2018), were performed in the same context (operant cages). In the present experiment, escalation of alcohol intake occurred in different cages than the maintenance, and the access (intermittent *vs*. daily), and concentrations of ethanol were also different between escalation procedure and the operant measure of consumption (20 *vs*. 10%). Thus, in this experiment, such differences between escalation and maintenance could have facilitated STN DBS effect on the maintenance of ethanol use in comparison with the cocaine experiment (Pelloux et al. 2018).

The level of consumption seems to be an important factor to ensure STN DBS efficiency since STN DBS decreased the consumption of alcohol only in HD rats. The lack of impact of STN DBS in LD rats might be attributed to a floor effect, implying that the LD rats did not consume enough ethanol to exhibit further a reduction triggered by STN DBS. Another hypothesis would be that the neurobiological effect of alcohol consumption does not induce the same changes in LD and HD rats. Indeed, when consuming alcohol, HD rats exhibit an increased power in gamma range and coherence in the medial prefrontal cortex (mPFC) compared to LD rats (Henricks et al. 2019). The mPFC is directly connected to the STN via the hyperdirect pathway (Maurice et al. 1998; Nambu et al. 2002). Additionally, abnormal increase in low frequency oscillations has been observed within the STN during cocaine escalation, and STN DBS was shown to reduce STN oscillatory activity (Pelloux et al. 2018). If such abnormal activity raised in the STN during alcohol escalation of HD rats, as that seen with cocaine, STN high frequency DBS could suppress it and also disturb the pathological oscillatory coherence between STN-mPFC as observed in PD (de Hemptinne et al. 2015; Delaville et al. 2015; Moran et al. 2012). This would lead to a decrease of alcohol intake only for HD rats after the loss of control.

Nonetheless, STN DBS appears to offer a potential therapeutic benefit for alcohol-related disorders, as suggested previously for other drugs.

#### …And of social influence during alcohol drinking

The influence of the social context was observed here in LD rats having escalated their alcohol consumption. Interestingly, in these animals, STN DBS effectively abolished the overconsumption induced by the presence of the peer. These results align with precedent findings, in which STN DBS has suppressed recreational alcohol overconsumption triggered by different types of peer (Sicre et al., unpublished data). STN optogenetic inhibition was also found to suppress the protective effect of a peer on cocaine recreational use (Vielle et al. preprint). The STN could thus be part of the neurobiological substrate of the influence of the social context on behavior. Indeed, studies focusing on the effects of ultrasonic vocalizations (USV) playback on cocaine intake showed that STN lesion abolished the beneficial influence of positive USV emitted by a stranger rat (Montanari et al. 2018; Vielle et al. 2021). Additionally, STN lesion suppressed the rewarding effect of positive USVs when rats were pressing a lever to listen to USVs (Vielle et al. 2021). It is important to note however that the combination of social context and STN manipulation does not systematically lead to neutralization or additivity of opposite effects, nor potentiation of similar effects. For example, STN lesions could decrease further the cocaine intake in presence of a peer (Giorla et al 2022), while STN neuronal optogenetic inhibition did not potentiate the decreased cocaine intake after escalation (Vielle et al preprint). All these findings suggest that presence of a peer effects on drug use probably involves STN as a neurobiological substrate. In human studies, STN DBS has been shown to induce deficits in facial and voice emotional decoding (Biseul et al. 2005; Brück, Kreifelts, and Wildgruber 2011; Kalampokini et al. 2020; Péron et al. 2010; 2013). A recent study has also shown that STN activity assessed with fMRI, was modulated by the presence of an observer during the performance of a stop-signal reaction time task by cocaine-addicted individuals (Terenzi et al. preprint). All these results converge to suggest the involvement of STN in mediating the impact of social interactions on drug-related behavior. Social interaction has been shown to trigger the release of oxytocin within the brain’s reward circuit (Dölen et al. 2013; Hung et al. 2017). Additionally, there are evidences suggesting that the STN possesses oxytocin receptors that can regulate locally dopamine transmission (Baracz and Cornish 2013, 2016). Furthermore the injection of oxytocin into the STN has been demonstrated to reduce methamphetamine-induced reinstatement (Baracz et al. 2015). Through oxytocin release induced by social interaction, the STN activity could thus be modulated and potentially affect drug consumption behavior.

Additional researches are required to elucidate further the role of the STN in social behaviors and its impact on drug-related effects.

## Acknowledgments

The authors thank Yann Pelloux for the custom-built interface and associated software, Joel Baurberg for technical assistance with all electronic devices, and the technical staff of the animal facility for ensuring the well-being of our animals. This work was funded by the Institut de Recherche en Santé Public and Alliance Aviesan in the call for proposal on substance use disorders and by the Institut de lutte contre le Cancer.

## Funding

Centre National de la Recherche Scientifique (CB, MD, NM)

Aix-Marseille Université (CB, CV, LV, MD, NM)

French Ministry of Higher Education, Research and Innovation (LV)

Institut de Recherche en Santé Public (IRESP-19-ADDICTIONS-02) (CB)

Institut National de lutte contre le Cancer (INCA_16032) (LV)

## Author contributions

Conceptualization: CB, CV, LV

Research performance: CV, LV, MW, NM

Data analyses: CV, LV

Funding acquisition: CB, LV

Writing – original draft: CB, CV, LV

Writing – review & editing: CB, CV, LV, MD, MW, NM

## Data and materials availability

All data are available in the main text, or on request to the authors.

